# Thalamostriatal and cerebellothalamic pathways in a songbird, the Bengalese finch

**DOI:** 10.1101/197590

**Authors:** David A. Nicholson, Todd Roberts, Samuel J. Sober

**Affiliations:** Graduate Program in Neuroscience, Atlanta, Georgia 30322; Department of Biology, Emory University, Atlanta, Georgia 30322; Department of Neuroscience, UT Southwestern Medical Center, Dallas, TX 75390-9111, USA

**Keywords:** thalamostriatal, songbird, basal ganglia, cerebellum

## Abstract

The thalamostriatal system is a major network in the mammalian brain, originating principally from the intralaminar nuclei of thalamus. Functions of the thalamostriatal system remain unclear, but a subset of these projections provides a pathway through which the cerebellum communicates with the basal ganglia. Both the cerebellum and basal ganglia play crucial roles in motor control. Although songbirds have yielded key insights into the neural basis of vocal learning, it is unknown whether a thalamostriatal system exists in the songbird brain. Thalamic nucleus DLM is an important part of the song system, the network of nuclei required for learning and producing song. DLM receives output from song system basal ganglia nucleus Area X and sits within dorsal thalamus, the proposed avian homolog of the mammalian intralaminar nuclei that also receives projections from the cerebellar nuclei. Using a viral vector that specifically labels presynaptic axon segments, we show in Bengalese finches that dorsal thalamus projects to Area X, the basal ganglia nucleus of the song system, and to surrounding medial striatum. To identify the sources of thalamic input to Area X, we map DLM and cerebellar-recipient dorsal thalamus (DT_CbN_). Surprisingly, we find both DLM and immediately-adjacent subregions of DT_CbN_ project to Area X. In contrast, a medial-ventral subregion of DT_CbN_ projects to medial striatum outside Area X. Our results suggest the basal ganglia in the song system, like the mammalian basal ganglia, integrate feedback from the thalamic region to which they project as well as thalamic regions that receive cerebellar output.

## Introduction

Motor skills learned by imitation and practice, like speaking a language or playing the piano, are under the control of a complex network of neural circuits. The basal ganglia and the cerebellum are key components of motor systems in the brain that contribute to learning and producing speech (Parrell, Agnew, Nagarajan, Houde, & Ivry, 2017; Watkins, 2011; Wolfram Ziegler, 2016; W Ziegler & Ackermann, 2017) and other complex motor skills (A. J. Bastian, 2006; Dudman & Krakauer, 2016; Manto et al., 2012; Shadmehr & Krakauer, 2008; Shmuelof & Krakauer, 2011). These contributions are thought to take place through basal ganglia and cerebellar output to motor thalamus, which in turn projects to cortex (Belyk & Brown, 2017; Bosch-Bouju, Hyland, & Parr-Brownlie, 2013; Ghez & Krakauer, 2000; Jürgens, 2002; Sommer, 2003; W Ziegler & Ackermann, 2017). However, in addition to projecting to cortex, thalamus projects to striatum. The thalamostriatal system is a major network in the mammalian brain that arises mainly from the intralaminar and midline nuclei (Smith et al., 2014). Output from the cerebellar nuclei can reach the basal ganglia through a subset of the thalamostriatal projections, providing a pathway through which the cerebellum can communicate with the basal ganglia (Chen, Fremont, Arteaga-Bracho, & Khodakhah, 2014; Hoshi, Tremblay, Feger, Carras, & Strick, 2005; Ichinohe, Mori, & Shoumura, 2000). However, it is unknown what if any role these pathways play in motor control.

Songbirds represent an ideal model system for investigating vocal learning and motor control, in part because of the structural and functional parallels between speech and birdsong (Bolhuis, Okanoya, & Scharff, 2010; Doupe & Kuhl, 1999; Marler, 1970). The songbird brain contains a network of discrete nuclei for learning and producing song, called the song system (Mooney, 2009), that shares many circuit-level and genetic characteristics with brain areas controlling speech (Konopka & Roberts, 2016). A thalamocortical-basal ganglia loop in the song system known as the anterior forebrain pathway (AFP, see Fig. 1) is required for juveniles to learn song and for adults to modify learned song (Andalman & Fee, 2009; Charlesworth, Tumer, Warren, & Brainard, 2011; Kao, Doupe, & Brainard, 2005). Although the AFP is thought to be homologous to similar loops in the mammalian brain, it is not known if songbirds have a thalamostriatal system similar to that of mammals. Specifically in the song system it is unknown whether in the AFP the basal ganglia nucleus, Area X, receives input from the thalamic nucleus DLM (Fig. 1, dashed dark blue line) or any other region of dorsal thalamus (Gale & Perkel, 2010). Dorsal thalamus is a strong candidate for the source of a thalamostriatal system in songbirds because it is the proposed avian homolog of the intralaminar nuclei in mammals (Veenman, Medina, & Reiner, 1997). Projections from other parts of dorsal thalamus to Area X could also provide a potential neuroanatomical pathway through which cerebellar output might reach the song system (Fig. 1, dashed red line and dashed light blue line) (Person, Gale, Farries, & Perkel, 2008). This would be particularly interesting given that it remains unknown whether the cerebellum is involved with learning and producing birdsong (Bolhuis et al., 2010; W Ziegler & Ackermann, 2017). We address these anatomical questions (Fig.1) in Bengalese finches (*Lonchura striata domestica*). This songbird species is of interest because it depends strongly on auditory feedback (Okanoya & Yamaguchi, 1997; S. M. Woolley & Rubel, 1997) and it has been shown to adapt its song to perturbations of auditory feedback (Sober & Brainard, 2009, 2012) in a manner reminiscent of cerebellar-dependent sensorimotor adaptation (Amy J Bastian, 2008; Parrell et al., 2017).

**Figure 1.**
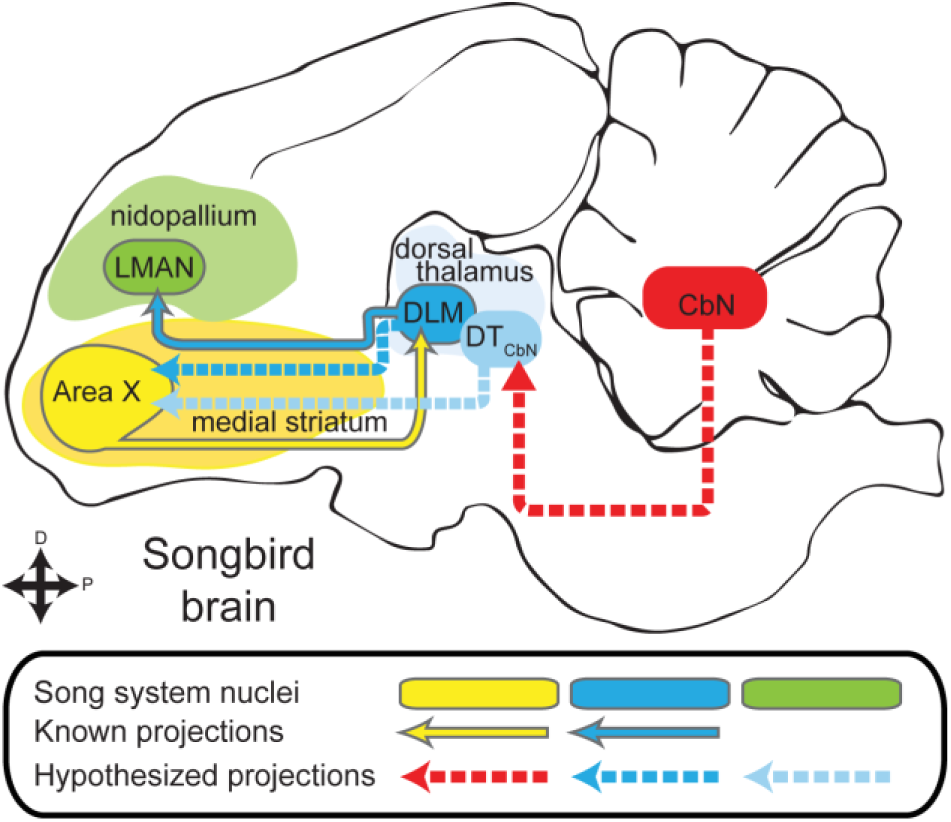
The anterior forebrain pathway (AFP) is a thalamocortical-basal ganglia loop in the song system required for learning song. It consists of basal ganglia nucleus Area X (yellow oval), thalamic nucleus DLM (blue oval) and cortical nucleus LMAN (green oval, known song system connections outlined in gray. The rest of the song system is not shown.). Using lentiviral tracing methods, we tested whether thalamus projects to the basal ganglia in a songbird, the Bengalese finch. Specifically we tested whether Area X (yellow oval) receives input from DLM (darker blue dashed arrow) or cerebellar-recipient dorsal thalamus (DT_CbN_, lighter blue dashed arrows). In order to identify targets of DT_CbN_, we first mapped out the projections of the cerebellar nuclei (CbN) to dorsal thalamus in detail (red arrow).

The question of whether a thalamostriatal system exists in songbirds has remained unanswered in part because of confounds that arise when using standard neuroanatomical tracers, as recognized previously (Bottjer, Halsema, Brown, & Miesner, 1989; Gale & Perkel, 2010; Person et al., 2008). Specifically, there is a passing fibers confound when retrograde tracers are used, because thalamocortical axons pass through the striatum. In addition, Area X in songbirds and the surrounding medial striatum (MSt) contain a specialized cell type that projects directly to thalamus, confounding results from standard tracers that travel in both the anterograde and retrograde directions. To avoid these confounds, we used lentiviral vectors that yield only anterograde label, and specifically label presynaptic axon segments. This method allowed us to show that dorsal thalamic neurons form synapses within the basal ganglia, including song system nucleus Area X. To determine which regions of dorsal thalamus project to the Area X, we use standard anatomical tracers to identify and map DLM as well as the regions of dorsal thalamus targeted by projections of the cerebellar nuclei (CbN). The latter region we refer to as cerebellar-recipient dorsal thalamus, DT_CbN_. We show that DLM and regions of DT_CbN_ surrounding DLM both project to Area X. More medial and ventral regions of DT_CbN,_ separate from DLM, project to medial striatum outside of Area X. Our findings suggest that thalamostriatal projections and cerebellar input to the basal ganglia may be general components of vocal motor control across vertebrates.

## Methods

All studies were carried out in adult (>90 days post hatch) male Bengalese finches either bought from a supplier or bred in our laboratory. Age of purchased birds was assessed by screening them for adult-like song. Work reported here was approved by the Emory University Institutional Animal Care and Use Committee.

### Surgery and tissue collection

We injected neuroanatomical tracers and lentiviral vectors with a stereotaxic apparatus (Leica/MyNeuroLab), using the co-ordinates in Table 1 to target brain regions described in the text. A summary of the injections and which figure they appear in is shown in Table 2. For all surgeries, we induced anesthesia with ketamine-midazolam and when necessary supplemented with isoflurane (0.25-2.5%). After induction, the bird’s scalp was anesthetized locally with ˜20uL lidocaine. An incision was made and the top layer of the skull was removed so we coutld localize the landmark “Y0”. We defined Y0 as the most posterior point visible at the junction of the midsagittal sinus and the two sinuses that run on either side of the cerebellum. After moving the pipette to the target co-ordinates on the surface of the skull, we made a craniotomy, opened the dura with a syringe needle, and then lowered the pipette to the target depth. Iontophoretic injections were made with an A&M 2100 stimulator, 7 seconds on, 7 seconds off, 4-10 µA, positive current, with a total injection time of 20-30 minutes. We used iontophoresis to inject fluorophore-tagged dextran amines, 10% in 0.1M phosphate buffer and adeno-associated virus expressing green fluorescent protein (AAV-GFP). For lentiviral vectors, we made pressure injections with a Nanoject II (Drummond). Lentiviral vectors were a 1:1 solution of mCherry and synapthophysin tagged with GFP. mCherry labeled cell bodies and axons, whereas synaptophysin-GFP specifically labeled synapses. These vectors were developed for anterograde tracing in songbirds; for details see (Bauer et al., 2008; Roberts, Klein, Kubke, Wild, & Mooney, 2008).

**Table 1.** Stereotaxic co-ordinates for targeting regions studied.

| Target | Anterior of y0 | Lateral of y0 | Depth | Beak bar angle below horizontal |
| --- | --- | --- | --- | --- |
| Dorsal thalamus | 0.9-1.1 | 1.3 | 4.1-4.3 | 45 |
| CbL | -1.2 – -1.6 | 1.3 – 1.4 | 3.25-3.45 | 50 |
| Area X | 5.5-5.7 | 0.9-2.2 | 2.9-3.1 | 20 |

**Table 2.** Injections for each experiment.

| Experiment | N | Figure |
| --- | --- | --- |
| **Test projection from thalamus to basal ganglia** |  |  |
| Injections of lentivirus in dorsal thalamus | 7* | 2,3,9,10 |
| **Test projection from CbN to dorsal thalamus** |  |  |
| Injections of dextran amines in CbL | 3 | 4,5 |
| Injections of dextran amines in dorsal thalamus | 3 | 6,7 |
| * 7 hemispheres from 6 birds |  |  |

Typical injection parameters for lentiviral vector injections with the Nanoject were 32.2 nl per press of the “inject” button, at a speed of 23 nl/second, for a total of 800-1500 nl in dorsal thalamus. Typically we waited 45 seconds between each press of the “inject” button, and left the pipette undisturbed for 5m after injecting the total volume, before raising slowly (˜100 µm every 5 seconds). After surgery, birds survived 5-7d when standard tracers were used, and 20-30d when lentiviral vectors were used. After the appropriate survival time, birds were sacrificed with an overdose of ketamine and midazolam, supplemented with isoflurane when necessary. The perfusate consisted of ˜20mL 0.95% saline with heparin, followed by ˜50mL 4% paraformaldehyde in 0.1M phosphate buffer (PB). The brain was removed and left in 4% paraformaldehyde, 30% sucrose solution overnight at room temperature, then transferred to 30% sucrose in 0.1M PB, where it was left until it sank in solution. Brains were cut parasagitally or coronally on a sliding freezing microtome and 30-60um sections were stored in 0.1M PB for further processing.

### Immunohistochemistry

Immunohistochemistical procedures were performed with antibodies at the dilutions given in Table 3, and according to the following procedure: after an initial rinse in 0.1M PB, we washed the tissue in 2% sodium borohydride in 0.1M PB for 0.5h, followed by three washes for 10m in 0.1M PB, as a form of epitope retrieval. Sections were then placed for 1h in a block solution of 2.5% normal donkey serum, 2.5% normal horse serum, 1% Triton-X 100 in 0.1M PB. Primary antibodies were diluted in 1% NDS, 1% NHS, 0.3% TX-100. Sections were incubated 24-48h at 4°C in primary solutions. Then the sections were rinsed 3x10m in 0.1M PB before incubating with secondary antibodies. Secondaries used with primaries to amplify lentiviral signal were tagged with fluorophores, while the secondary used with the parvalbumin primary was biotinylated so it could be further incubated with Vector labs streptavidin-AMCA (SA-5008) as a tertiary. Both secondary and tertiary solutions consisted of 0.3% TX-100 in 0.1M PB, and incubations in these solutions lasted 1h. In cases where a tertiary was used, the incubation was preceded by 3x10m washes in 0.1m PB. In all cases, sections were washed 3 more times for 10m in 0.1M PB before they were mounted on Fischer Superplus slides, then left overnight to dry. Sections were briefly rehydrated before using Fluoro-Gel with DABCO mounting medium to apply coverslips that were then sealed with clear coat fingernail polish.

**Table 3.** Antibodies used.

| **Name** | **Immunogen structure** | **Manufacturer, catalog #, RRID, species, mono/poly** | **Concentration** |
| --- | --- | --- | --- |
| Anti-red fluorescent protein | mushroom polyp coral Discosoma | Rockland 600-401-379, AB_2209751, Rabbit polyclonal | 1:1000 |
| Anti-mCherry | full-length protein mCherry | Life technologies/Invitrogen M11217, AB_2536611, Rat monoclonal | 1:1000 |
| GFP tag (clone 3E6) | GFP isolated directly from Aequorea victoria | Life technologies/Invitrogen A-11120, AB_2534132, Mouse monoclonal | 1:2000 |
| GFP tag | GFP isolated directly from Aequorea victoria | Life technologies/Invitrogen A-11122, AB_2534134, Rabbit polyclonal | 1:2000 |
| Parvalbumin | Parvalbumin purified from frog muscle | EMD Millipore MAB1572, AB_2174013, Mouse monoclonal | 1:2000 |
| Rhodamine Red X donkey anti rabbit | whole molecule rabbit IgG | Jackson 711-295-152, AB_2340613, donkey polyclonal | 1:400 |
| Rhodamine Red X donkey anti rat | whole molecule rat IgG | Jackson 712-295-150, AB_2340675, donkey polyclonal | 1:400 |
| Alexa 488 donkey anti-mouse | whole molecule mouse IgG | Jackson 715-545-150, AB_2340846, donkey polyclonal | 1:400 |
| Alexa 488 donkey anti-rabbit | whole molecule rabbit IgG | Jackson 711-545-152, AB_2313584, donkey polyclonal | 1:400 |
| Biotinylated horse anti-mouse IgG (H+L) | whole molecule mouse IgG | Vector laboratories BA-2000, AB_2313581, horse, polyclonal | 1:400 |

In initial experiments (Fig. 2) we used the Rockland rabbit anti-RFP and Invitrogen mouse anti-GFP antibodies to detect mCherry and GFP-synaptophysin expression. In later experiments where we used the Millipore mouse anti-Parvalbumin as a marker for Area X and other song system nuclei, we could not use the mouse anti-GFP and so instead we used the Invitrogen rat anti-mCherry and the Invitrogen rabbit anti-GFP.

**Figure 2.**
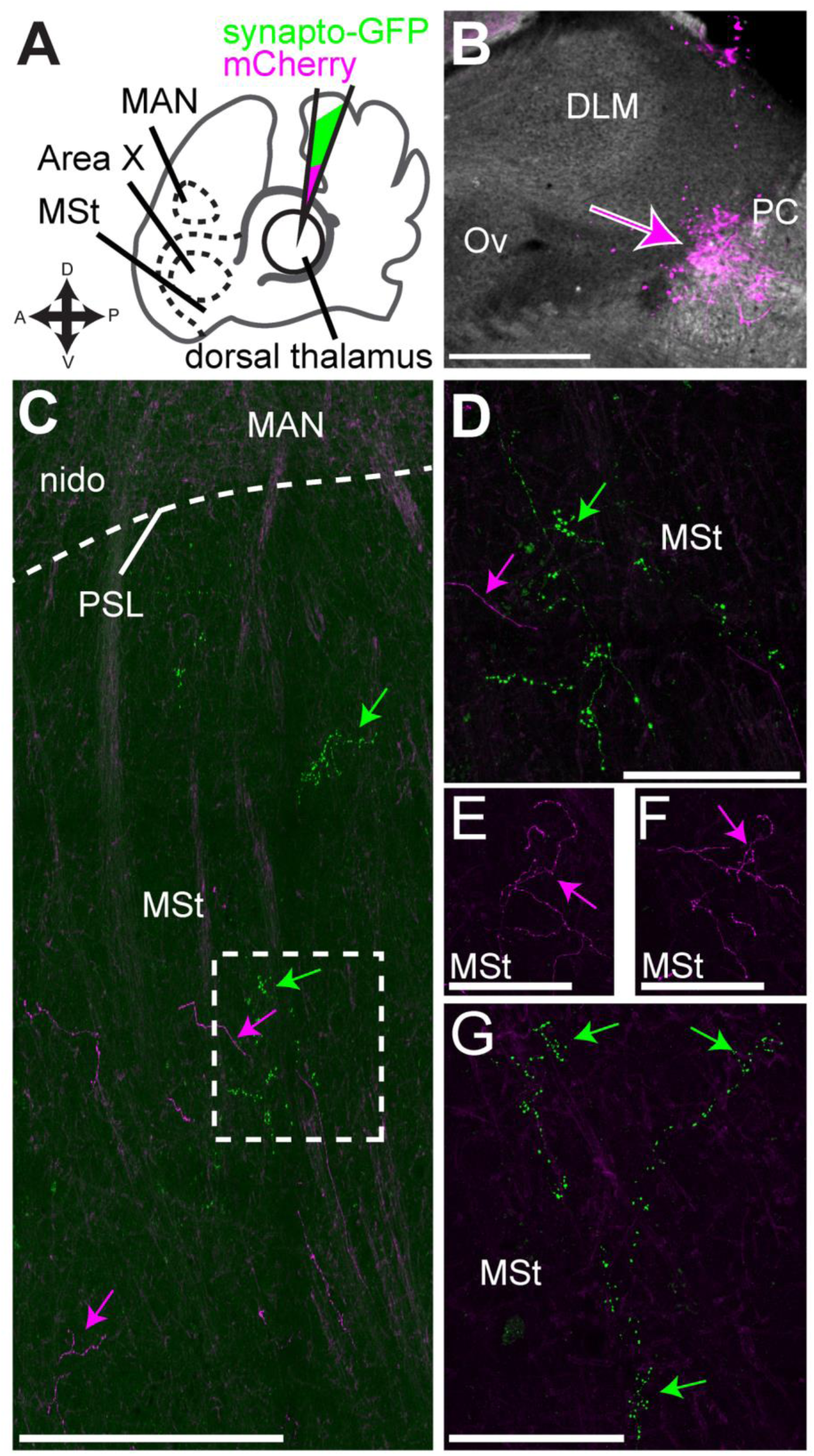
Dorsal thalamus projects to striatum **A,** Schematic of experiment. We injected into dorsal thalamus (DT) a 1:1 solution of two lentiviral vectors, one expressing synaptophysin-GFP and the other mCherry, and then looked for label in medial striatum (MSt) which contains basal ganglia nucleus of the song system, Area X. One landmark we used to determine whether sections included MSt was the magnocellular nucleus of the anterior nidopallium (MAN) which is obvious even in unstained tissue. **B,** Widefield image of injection site from representative case. This injection was outside of thalamic nucleus of the song system DLM, just anterior to the posterior commissure (PC) and in the same plane as auditory thalamus nucleus ovoidalis (Ov) **C,** Image showing mCherry and synaptophysin-GFP labeled processes in MSt ventral to MAN. Magenta arrows, mCherry signal, green arrows, synaptophysin-GFP signal. PSL, pallial-subpallial lamina, nido, nidopallium outside of MAN. **D,** Synaptophysin-GFP labeled processes in MSt had varicosities, suggesting they were synapses. Higher-power scan of the area outlined in **C** with a dashed box. **E, F.** mCherry labeled processes could also be found in MSt with varicosities, indicating this is the actual morphology of the axons and did not result from ectopic (over)expression of synaptophysin-GFP **G.** Another example of synaptophysin-GFP labeled processes in MSt. **E-G** are all in the same section within 500 µm of each other but are enlarged to be easily visible. Scale bars: **B-C,** 500 µm; **D-G,** 100 µm.

### Production and specificity of antisera

Please see Table 3 for list of antibodies used.

Anti-red fluorescent protein (RFP) antibody, Rockland 600-401-379, AB_2209751, Rabbit polyclonal, detects RFP but not GFP as shown by Western blot (manufacturer’s datasheet). The antibody has been used previously to amplify mCherry signal expression from viral vectors (De Arcangelis, Liu, Soto, & Xiang, 2009; Dinh & Bernhardt, 2011) including vectors used in neuroscience studies (Redondo et al., 2014; Sreenivasan, Karmakar, Rijli, & Petersen, 2015).

Anti-mCherry, Life technologies/Invitrogen M11217, Rat monoclonal, specifically detects mCherry as shown with Western blot and flow cytometry (manufacturer’s datasheet). The antibody has been used previously in virally-mediated neural tracing experiments (Schwarz et al., 2015).

Anti-GFP, Life technologies/Invitrogen A-11120, mouse monoclonal, was raised against GFP isolated from Aequorea Victoria (manufacturer’s datasheet). It has been shown to detect GFP fusion proteins expressed in neurons under genetic control (Busch, Selcho, Ito, & Tanimoto, 2009; Liu, Luo, Carlsson, & Nässel, 2015) and GFP expression resulting from viral vectors (Keen-Rhinehart et al., 2009; Vujovic, Gooley, Jhou, & Saper, 2015)

Anti-GFP, Life technologies/Invitrogen A-11122, rabbit polyclonal, was raised against GFP isolated from Aequorea Victoria (manufacturer’s datasheet). Previous reports show this antibody detects GFP expression induced in neurons by viral vectors (Davis et al., 2011; Lindberg, Chen, & Li, 2013)

Anti-parvalbumin, EMD Millipore MAB1572, mouse monoclonal, was raised against parvalbumin purified from frog muscle, and in Western blots yields a band at 12kDa (the weight of parvalbumin). The antibody shows specific immunoreactivity with parvalbumin expressing interneurons (McKenna et al., 2013) and has been used to label such neurons in songbirds (Li et al., 2013). It has previously been shown that one cell type in Area X expresses parvalbumin and that Area X shows higher immunoreactivity for parvalbumin in its neuropil than the surrounding medial striatum (Braun, Scheich, Schachner, & Heizmann, 1985; Carrillo & Doupe, 2004; Reiner, Laverghetta, Meade, Cuthbertson, & Bottjer, 2004)

### Microscopy, digital photography, and image processing

Low-power widefield images were obtained with a Zeiss Axioplan 2 and Olympus IX51. Confocal z-stacks were obtained with Leica SP8 inverted and Olympus FV1000 inverted microscopes. Brightness and contrast were adjusted using ImageJ, Zeiss AxioVision software (for images acquired with the Axioplan 2), or Photoshop (for images acquired with the Olympus IX51). ImageJ was used for all processing of z-stacks, including z projections, adjustment of brightness and threshold, and changing of look-up tables (e.g., to convert red to magenta). We used the following procedure to calculate the distances from the midline shown in figures on parasagittal sections: we chose the zero point to be the middle of the interstitial nucleus of Cajal (InC), which is found at the midline and which we observed usually occurred in two sections in parasagittal sections; we then counted the number of sections including the section in the figure and the section of InC between that section and the zero point; we multiplied that number of sections by the section thickness (e.g. 20 sections x 40 micrometers/section) and lastly we multiplied the total number of micrometers by 1.5, a factor to account for shrinkage that occurred when the brain was fixed. We found this conversion factor by measuring the distance between injection sites and the midline in fixed tissue and solving for the average value that would convert this distance back to the mediolateral distance we used for stereotaxic targeting the injections.

### “Drawings” of signal from lentiviral injections

To present results from lentiviral injections, we followed a procedure that yielded figures similar to camera lucida-assisted drawings of light microscopy material. After performing fluorescence immunohistochemistry on sections to amplify signal, we made large tiled scans of the sections with a confocal microscope using a 40x objective. We then made z-projections of these tiled scans, compressing the z-stack into one x-y plane, and adjusted the brightness and contrast in ImageJ. In Adobe Illustrator, we aligned the tiled z-projection of each section with a widefield image taken with a 4x objective of the same section. The 4x objective was used with a DAPI filter to image the Parvalbumin (PV) signal that allowed us to identify the borders of Area X and other areas of interest. To ensure that the 40x images and the 4x images were at the same scale, we placed scale bars of the same size on both images and aligned the scale bars before aligning the images. We then made the 40x images transparent in Illustrator and aligned the actual sections by eye using landmarks, e.g., the edges of the sections and cytoarchitectural landmarks that were visible because of slight background autofluorescence. Using a Wacom graphics tablet with a stylus, we outlined regions of interest like Area X based on the PV signal. Next we imported the aligned images and the outlines of regions into Adobe Photoshop as separate layers. On the layers with the 40x tiled z-projections, we used the Lasso tool to outline all areas of signal (either synaptophysin-GFP imaged with Alexa 488 secondaries or mCherry imaged with Rhodamine Red X). We copied these areas with signal to a separate layer and then applied the Invert and Threshold functions. Next we opened the files again in Illustrator and used the Image Trace tool on the inverted and thresholded signal (mode: black and white, threshold: 210-245), paths: 100%, corners: 75%, noise: 15-25px, create: fills, snap curves to lines: no, ignore white: yes). Lastly we selected “Make and Expand” to convert the Image Trace objects to vector art, and then colored the vectors, either magenta for the mCherry signal or green for the synaptophysin-GFP signal. We exported the completed tracing as a .png file.

Simply because of space constraints, we show only representative sections from the series of tracings in Figs. 9 and 10. The entire series of tracings is included in the Supporting Information (<u>https://doi.org/10.6084/m9.figshare.5437975</u>).

### Map of dorsal thalamus

To determine whether song system thalamic nucleus DLM or cerebellar-recipient dorsal thalamus (DT_CbN_) were the source of projections to Area X or the surrounding medial striatum, we carefully mapped both regions of dorsal thalamus (Figs. 4, 5, and 8). Having done this, we could then superimpose sites of injection into dorsal thalamus on this map and see whether an injection that had produced synaptophysin-GFP label in Area X had been in DLM or DT_CbN_. As we show, the projections of CbL were consistent across animals (Fig. 5). Likewise we saw that DLM as defined by its input from Area X was consistent across animals (Fig. 8). In addition we could easily recognize DLM even in unstained tissue because of the heavier myelination that showed up as a brighter area when sections were viewed with darkfield through a DIC filter of a 5x objective (Fig. 8). To produce a reference map of dorsal thalamus, we outlined areas in one bird where we injected a GFP expressing viral vector in Area X to label its projection to DLM, and injected dextran amines in CbL to label DT_CbN_. We used only one bird as a reference so that we could be sure there was no added noise due to slight differences in alignment when sectioning brains and then aligning series of section by eye. Hence in this reference series we could confidently localize DLM and DT_CbN_ in each section relative to each other.

We aligned this map with the injection sites from viral injections in dorsal thalamus by using cytoarchitectural landmarks that were clearly visible when viewing unstained sections at 5x with darkfield. (We combined DIC with darkfield simply because this increased contrast on the microscope used to image injection sites, a widefield Zeiss Axioplan 2). Typical injection sites consisted of neurons expressing either synaptophysin-GFP or mCherry, with a sparse population of cells expressing both. We chose to define injection sites by the mCherry labeled cell bodies because the mCherry label was easier to image at 5x. Comparison of this label with synapthophysin-GFP label in the injection site, as imaged with a confocal, showed no obvious difference between the injection site as defined by mCherry signal or synaptophysin-GFP signal—i.e. injection sites appeared to be mostly homogenous mixture of neurons infected with one or the other viral vector.

Results from the bird that we used as a reference for the map of DLM and DT_CbN_, as well as raw images of the injection sites from each case in Fig. 9 and 10, and the files showing how they were aligned with this map, are included in the Supporting Information (<u>https://doi.org/10.6084/m9.figshare.5437975</u>).

## Results

### Dorsal thalamus projects to medial striatum, including Area X

To determine whether dorsal thalamus projects to medial striatum, we used lentiviral vectors for neuroanatomical tracing. There were two advantages of using these lentiviral vectors. The first is that they produce only anterograde label, because they infect only cell bodies at the injection site (Grinevich, Brecht, & Osten, 2005; Roberts et al., 2008) whereas standard tracers are picked up by the cell body and by axon terminals, traveling to some extent in both the anterograde and retrograde direction (Kobbert et al., 2000; Reiner et al., 2000). If we had used standard tracers, we would not have been able to distinguish between anterograde and retrograde label in Area X, because both Area X and the surrounding medial striatum project directly to dorsal thalamus. The second advantage of the vectors we used is that one contained GFP-tagged synaptophysin, allowing us to specifically label presynaptic axon segments with GFP signal (synaptophysin protein is localized at the presynaptic axon terminal and axon segments proximal to the terminal (Grinevich et al., 2005; Roberts et al., 2008)).

We made an initial series of large injections in dorsal thalamus to simply test whether any part of dorsal thalamus projected to any part of the medial striatum (including Area X). These injections (experiment schematic, Fig. 2A) contained a 1:1 ratio of the synaptophysin-GFP vector and another expressing mCherry (Roberts et al., 2008). The mCherry vector labels cell bodies and axons, allowing us to identify the injection site in dorsal thalamus (Fig. 2B). In no case did we see retrograde label (e.g., label of cell bodies in Area X, medial striatum, or CbL) from injections in dorsal thalamus, making us confident that our results are based exclusively on signal produced by infection of cell bodies local to the injection site and transported in the anterograde direction. We found that injections in dorsal thalamus yielded synaptophysin-GFP label throughout the medial striatum (MSt, Fig. 2C, green arrows). As stated in the introduction, the basal ganglia nucleus of the song system, Area X, sits within MSt of songbirds. We suspected that anterograde label was also present in Area X, based on landmarks, such as the presence of cortical nucleus LMAN within a section. We investigate this using immunohistochemistry to identify the borders of Area X in the next set of experiments described. The axon segments labeled by synaptophysin-GFP had numerous varicosities (Fig. 2D), suggesting they formed synapses *en passant*, as seen in the mammalian thalamostriatal system (Berendse & Groenewegen, 1990; Deschenes, Bourassa, & Parent, 1996). We also saw mCherry labeled processes with varicosities (Fig. 2E, F), indicating that these varicosities did not arise as a result of overexpression of synaptophysin. Qualitatively, it appeared that the synaptophysin-GFP (as opposed to mCherry) preferentially labeled the processes that form varicosities within the medial striatum (Fig. 2G, green arrows).

Having shown that dorsal thalamus projects to the basal ganglia, we then sought to demonstrate conclusively whether thalamostriatal projections specifically target Area X. We made another series of viral injections in dorsal thalamus (Fig. 3A, B), and for each case we also labeled Area X by performing immunohistochemistry against parvalbumin (PV) (Fig. 3B, C). Neuropil in Area X is more strongly enriched in PV than the surrounding medial striatum (Reiner et al., 2004). This allowed us to clearly identify Area X in the each section and determine whether processes labeled by synaptophysin-GFP and mCherry were within it. By aligning the PV-stained section with mosaic images of high-powered confocal stacks imaging the GFP and mCherry, we could see that some part of dorsal thalamus projected to Area X (Fig. 3D). These processes again had varicosities characteristic of thalamostriatal axon terminals (Fig. 3E)

**Figure 3.**
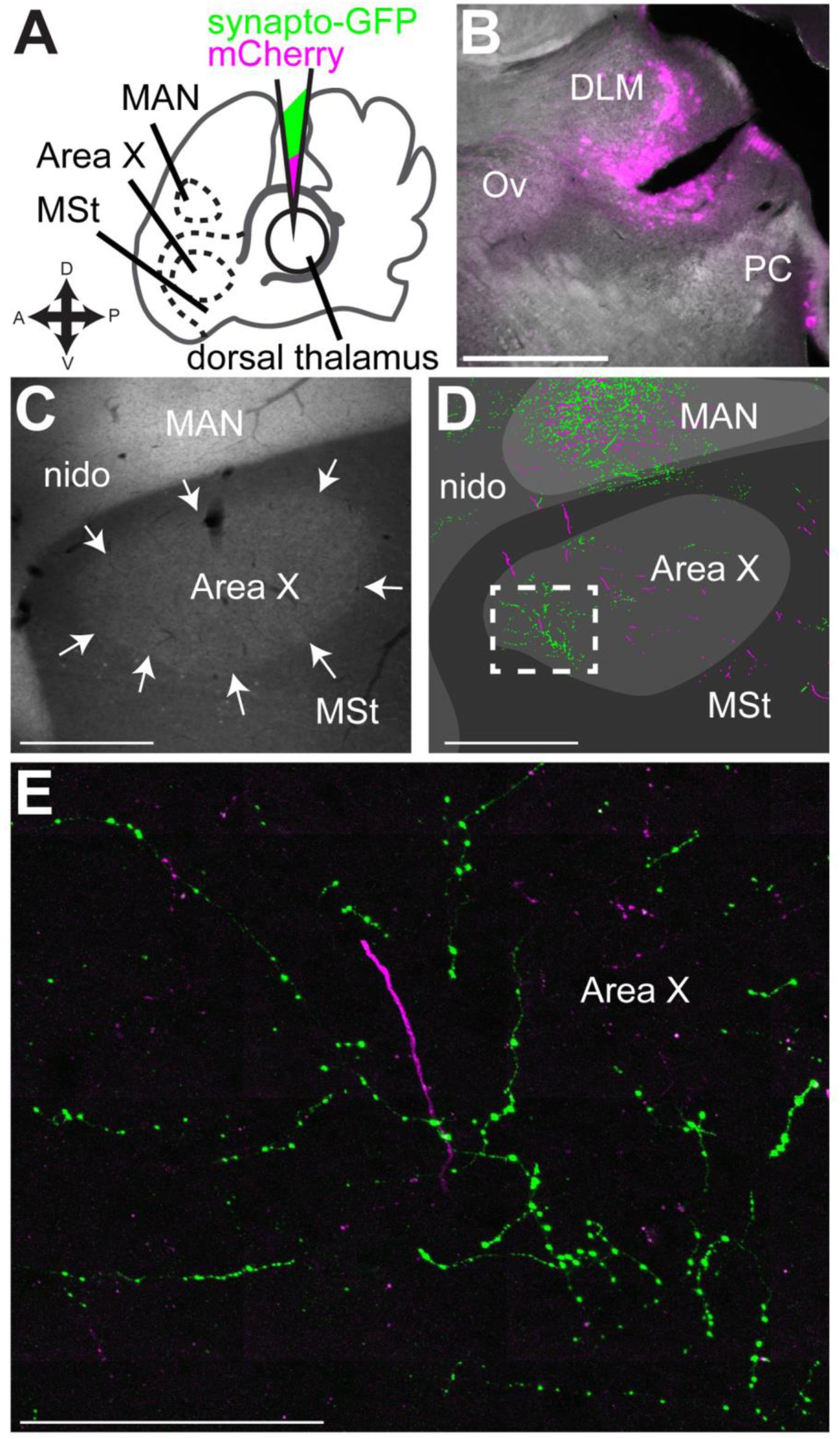
Dorsal thalamus projects to striatum, including Area X. **A,** Schematic of experiment. As in Fig.2, we injected into dorsal thalamus a 1:1 solution of two lentiviral vectors, one expressing synaptophysin-GFP and the other mCherry, but in these experiments we labeled Area X immunohistochemically with antibodies against parvalbumin, and then determined whether viral label was found in Area X or medial striatum (MSt) as well as the magnocellular nucleus of the anterior nidopallium (MAN). **B,** Widefield image of injection site from representative bird. **C,** Parvalbumin stain to outline Area X (white arrows) from the same bird. **D,** “Camera lucida” style tracing of GFP and mCherry signal from same section shown in (c). **E,** Confocal image of area shown in white box in **D**. Note varicosities suggesting *en passant* synapses. Scale bars in **A-D**, 500 µm. Scale bar e in e, 250 µm.

## The cerebellar nuclei project to contralateral dorsal thalamus

Although our results showed that dorsal thalamus projects to Area X and the surrounding medial striatum, they did not clearly demonstrate which regions of dorsal thalamus gave rise to the projections to Area X. Based on previous work, we hypothesized that the source of projections to Area X would be either DLM or cerebellar-recipient dorsal thalamus (DT_CbN_). To determine whether DLM or DT_CbN_ project to Area X or medial striatum, we first mapped these areas of dorsal thalamus using standard tracers and viral vectors.

We began by identifying the region of dorsal thalamus targeted by the cerebellar nuclei in Bengalese finches. Our initial set of injections targeted the lateral cerebellar nuclei (CbL) because of the previous work in other songbird species suggesting this was the main source of cerebellar projections to thalamus (Person et al., 2008; Vates, Vicario, & Nottebohm, 1997). Injections of dextran amines in CbL yielded anterograde label across the mediolateral extent of contralateral dorsal thalamus (Fig. 4). This result was consistent across animals (n=3, Fig. 5). In every case where we successfully targeted CbL, we saw label across the entire mediolateral extent of dorsal thalamus, from near the midline (Fig. 4K) to very laterally near the pretectal nucleus (Pt) (Fig. 4C). We always saw a densely labeled area posterior to song system thalamic nucleus DLM (Fig. 5C, black arrows with white outline), in roughly the same mediolateral plane as the auditory region of thalamus, nucleus ovoidalis (Ov). (Results confirming the location of DLM in relation to DT_CbN_ are described in the next section.) Cerebellar axon terminals surrounding but not invading DLM were also seen after injections in the cerebellar nuclei (CbN) in zebra finches (Person et al., 2008). The area of dense label extended laterally to the same mediolateral plane as the retinal-recipient nucleus DLL (Fig. 5D, white arrows indicate label from cerebellar injection), posterior to the region that receives retinal output (H. J. Karten et al., 2013). We note that in two of three cases shown in Fig. 5, injections in CbL also yielded some retrograde label of the medial spiriform nucleus (SpM). SpM is known to project to the cerebellum (H. Karten & Finger, 1976; Person et al., 2008), specifically the cerebellar cortex (J. M. Wild, 1992). In one of the three cases, SpM was retrogradely labeled on both sides of the brain, and on the ipsilateral side where there was no anterograde label from CbL, we saw label of axon collaterals in dorsal thalamus, i.e., it appeared that the tracer traveled retrogradely from the cerebellum to SpM cell bodies and then from SpM traveled anterogradely to label collaterals in dorsal thalamus. However there were few of these collaterals and the amount of signal was very sparse compared to the strong anterograde label seen from all CbL injections (the sparse label of SpM collaterals is shown in the Supporting Information, <u>https://doi.org/10.6084/m9.figshare.5437975</u>). We are therefore confident that the majority of label in contralateral dorsal thalamus in all cases traveled anterograde from CbL. In addition to the label in dorsal thalamus, we also saw widespread signal throughout the midbrain, with dense innervation of the red nucleus, ansa lenticularis, and SpM. These results are consistent with what was reported by Person et al. (2008) and what is seen in other bird species (Arends & Zeigler, 1991) and mammals (Hoshi et al., 2005; Loreta Medina, Veenman, & Reiner, 1997). Since the objective of these experiments was to map the cerebellar-recipient regions of dorsal thalamus, we do not describe these other targets of the cerebellar nuclei further, but we do include the results in the Supporting Information (<u>https://doi.org/10.6084/m9.figshare.5437975</u>). In the Discussion we address how other targets of CbL in the midbrain may relate to the function of the cerebellothalamic projections.

**Figure 4.**
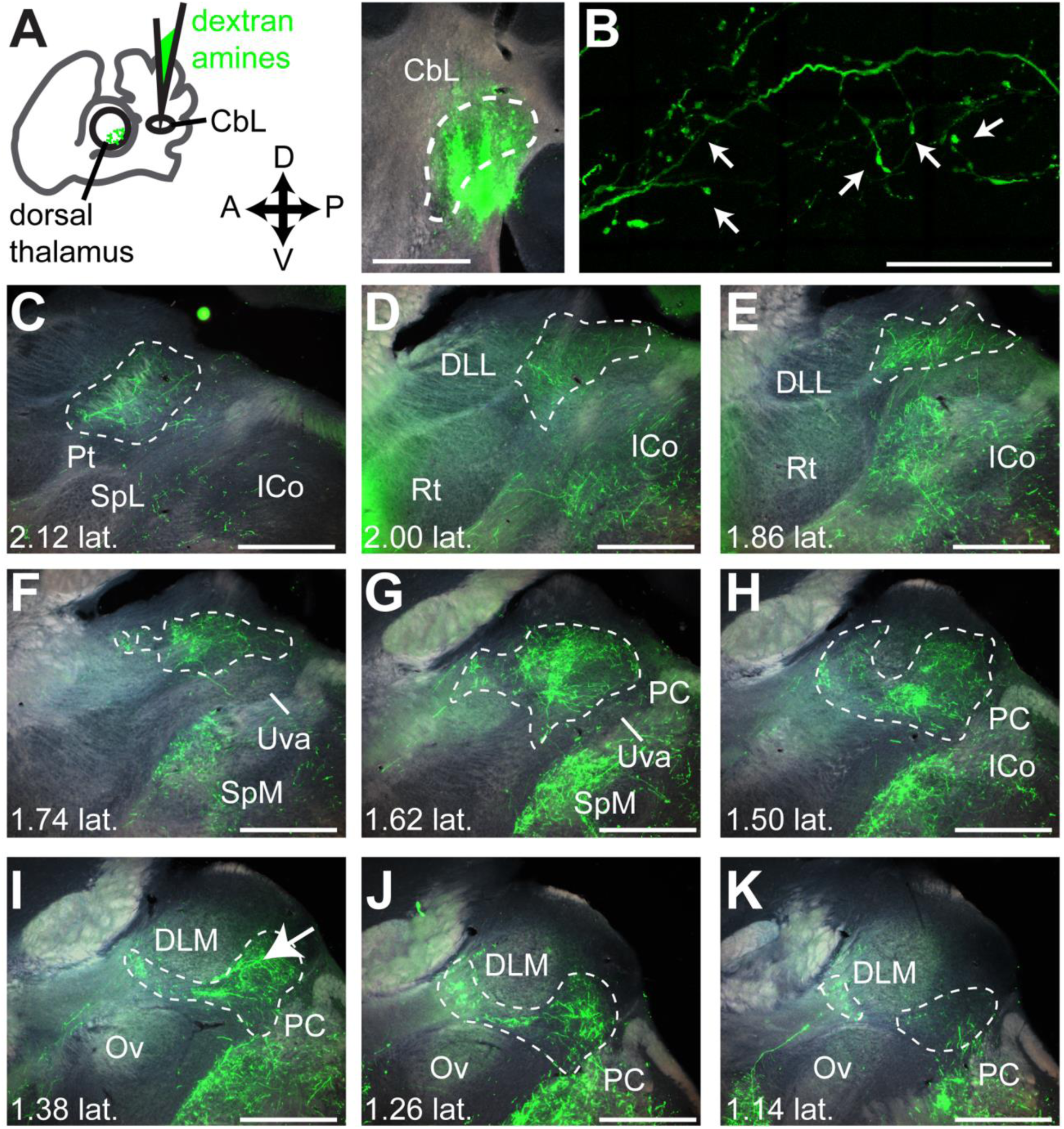
The lateral cerebellar nucleus (CbL) projects to contralateral dorsal thalamus. **A,** Schematic (left panel) of experiment showing injection in CbL and widefield image (right) showing injection site. **B,** High-resolution confocal image of axon-terminal like morphology in contralateral dorsal thalamus. This image was taken from the area indicated with a white arrow in **I**. **C-K**, representative series of widefield images across dorsal thalamus showing anterograde label from the injection in CbL. **C** is the most lateral section and **K** is the most medial. Estimate of distance from midline (e.g. “1.02 lat.”) is given in millimeters, calculated as explained in Methods. Dotted white lines demarcate the areas we considered cerebellar-recipient dorsal thalamus. Left is anterior and up is dorsal. This series is from “bird 3” in Figure 5. All scale bars 500 µm. Abbreviations from C-K: Pt, pretectal nucleus. SpL, lateral spiriform nucleus. ICo, inferior colliculus. DLL, dorsolateral thalamus, lateral part. Rt, nucleus rotundus. Uva, uvaeform nucleus. SpM, medial spiriform nucleus. PC, posterior commissure. Ov, nucleus ovoidalis. DLM, dorsolateral thalamus, medial part.

**Figure 5.**
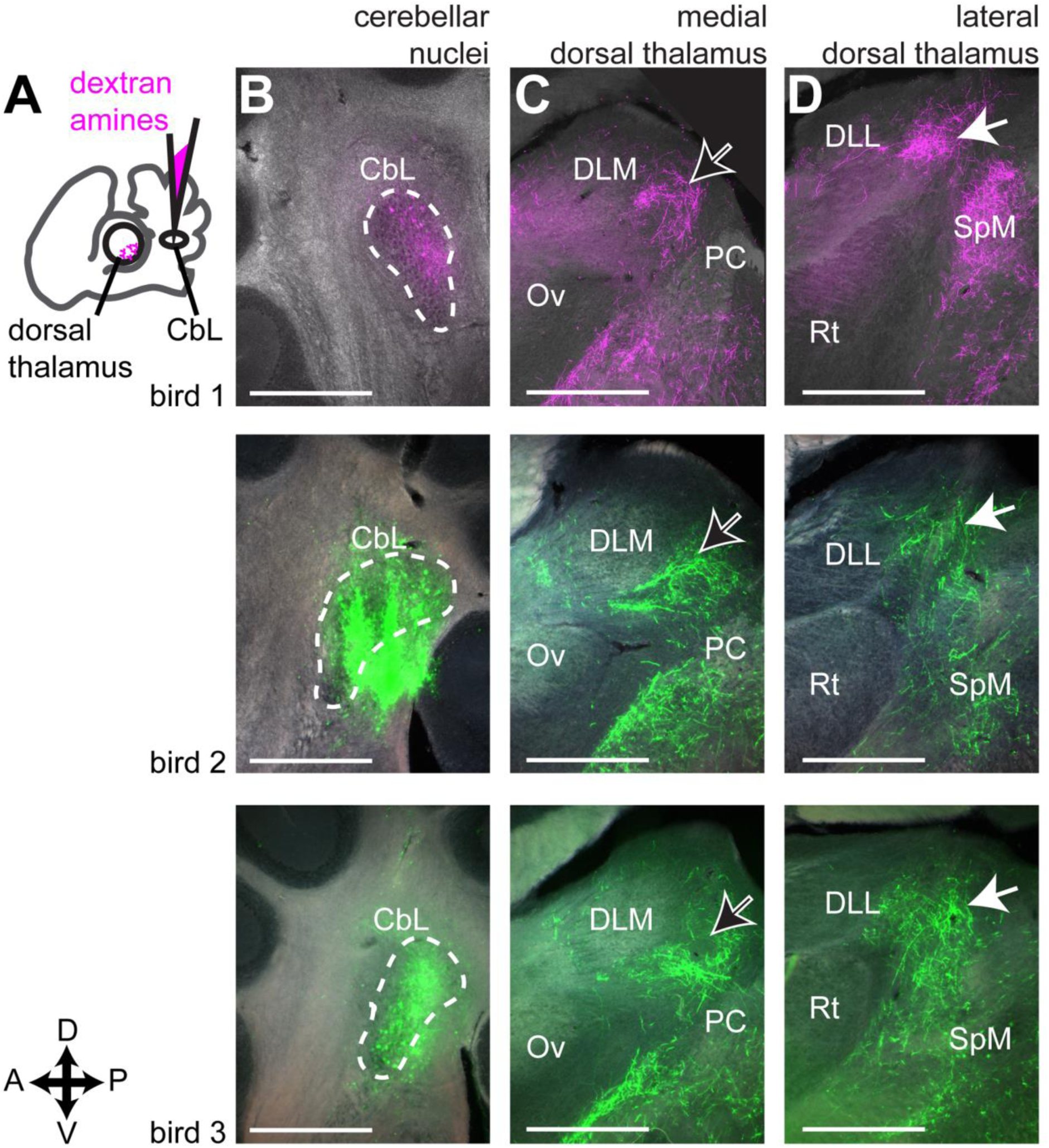
CbL axon terminals target medial dorsal thalamus adjacent to song system nucleus DLM, but also target more lateral dorsal thalamus. **A,** Schematic representation of experiment, showing injection site in lateral cerebellar nuclei, and anterograde label in contralateral dorsal thalamus. **B,** Injection sites. In one case shown (top row), the fluorophore conjugated to the dextran amines was tetramethylrhodamine. In the other cases, the conjugate was fluorescein. The choice of fluorophore did not affect results. **C,** More medial site in dorsal thalamus with strong label. DLM, dorsolateral thalamus, medial part. Ov, nucleus ovoidalis. PC, posterior commissure. **D,** More lateral site in dorsal thalamus with strong label. DLL, dorsolateral thalamus, lateral part. Rt, nucleus rotundus. SpM, medial spiriform nucleus. All sections are parasagittal, left is anterior and up is dorsal. All scale bars 500 µm.

To confirm that CbL projects to dorsal thalamus, we made injections of dextran amines in dorsal thalamus and looked for retrograde label in CbL (Fig. 6A). These injections yielded retrograde label of contralateral CbN (Fig. 6B, 6C). As previously reported for songbirds (Vates et al., 1997), the injections gave strong label in CbL. In addition, we saw retrograde label in intermediate and medial regions of the cerebellar nuclei (CbI and CbM, Fig. 6D), consistent with what has been reported for other bird species (Loreta Medina et al., 1997) and for mammals (Hoshi et al., 2005). We combined results from animals (n=3) in which the mediolateral position of the thalamic injection site varied (Fig. 7) to map of the regions of cerebellar nuclei that project to dorsal thalamus in a songbird. In two of these birds we made multiple injections in the mediolateral plane of dorsal thalamus (Fig. 7, injection sites colored cyan and magenta), which yielded strong retrograde label. A more ventral and medial injection (Fig. 7, injection site colored yellow) yielded less retrograde label. All three injections yielded the strongest retrograde label in CbL, but they also yielded label of many neurons in CbI and a few neurons in CbM as well (Fig. 7B). Hence our results suggest that the strongest projection to dorsal thalamus originates from CbL, but that there is a significant contribution from CbI, and some input from CbM as well.

**Figure 6.**
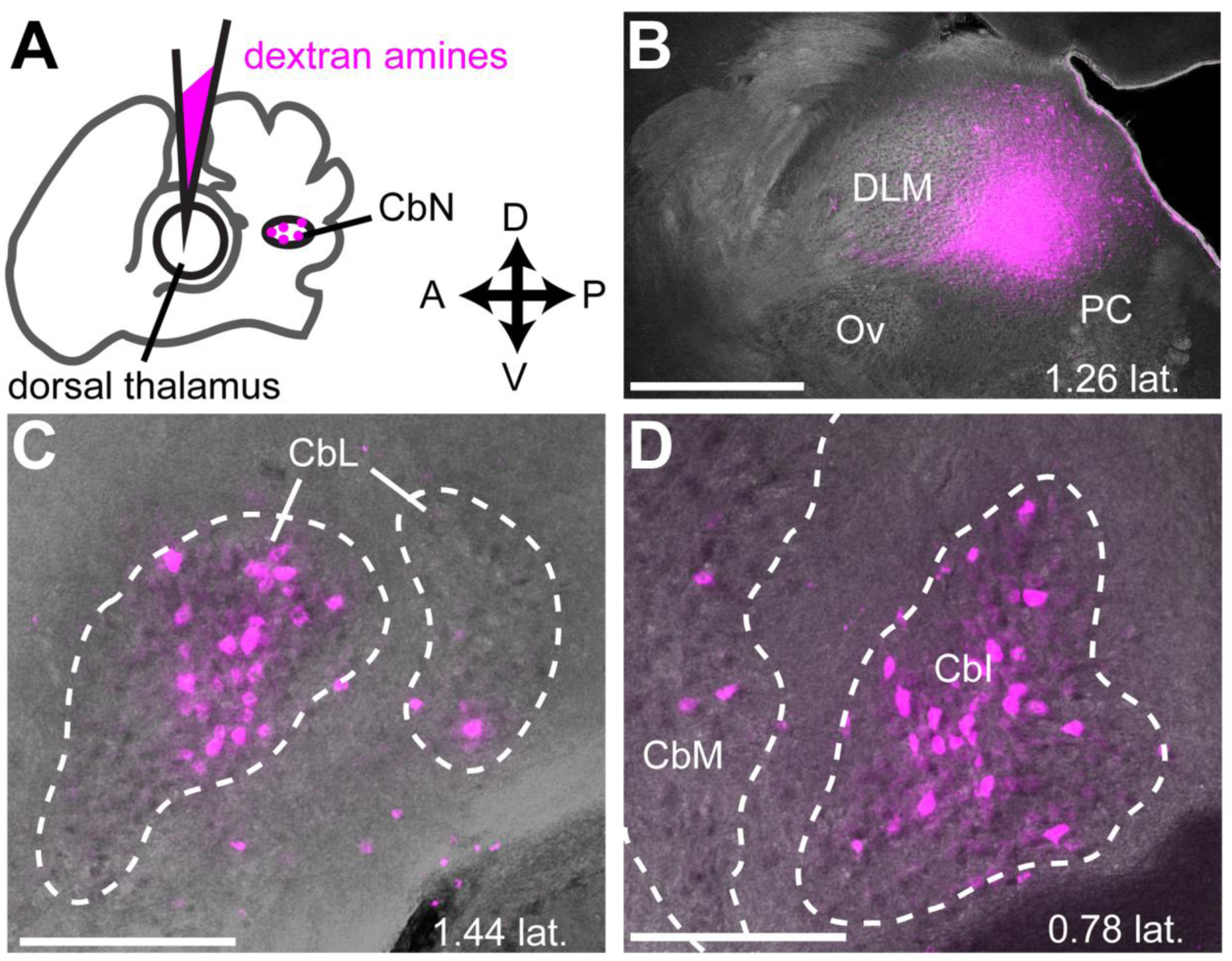
Injections in dorsal thalamus yield retrograde label in the cerebellar nuclei. **A,** schematic indicating injection site in dorsal thalamus and site of retrograde label in contralateral cerebellar nuclei (CbN) (shown in same plane of cartoon “section”). **B,** Representative injection site in dorsal thalamus. Parasagittal section. Anterior is left and dorsal is up. DLM, dorsolateral thalamus, medial part. Ov, nucleus ovoidalis. PC, posterior commissure. **C,** Retrograde label in contralateral lateral cerebellar nucleus (CbL). **D,** Retrograde label in contralateral intermediate (CbI) and medial (CbM) cerebellar nucleus. Scale bar, 500 µm

**Figure 7.**
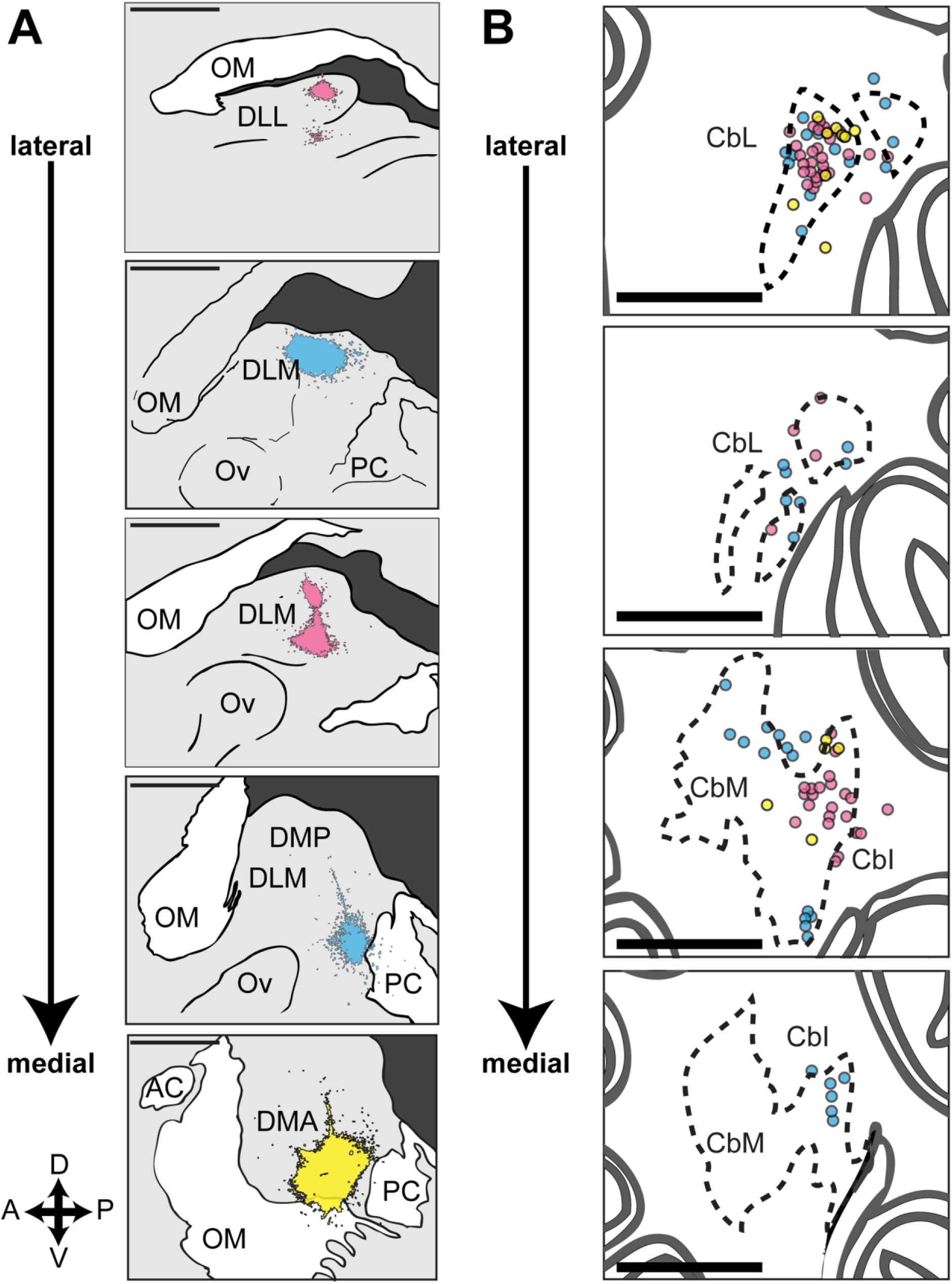
All of the cerebellar nuclei project to dorsal thalamus. **A,** Injection sites in dorsal thalamus from 3 birds (magenta, cyan, yellow) arranged from lateral to medial. Note that the 3rd panel from the top corresponds approximately to the injection site shown in figure 5A. DLL, dorsolateral thalamus, lateral part. OM, occipito-mesencephalic tract. DLM, dorsolateral thalamus, medial part. Ov, nucleus ovoidalis. PC, posterior commissure. **B,** Schematic of retrograde label in the contralateral cerebellar nuclei arranged from lateral to medial.n Colored circles represent retrogradely filled cell bodies. Color of each circle indicates retrograde label from injection site with the same color in **A**. CbL, lateral cerebellar nucleus. **D,** CbI, intermediate cerebellar nucleus. CbM, medial cerebellar nucleus. All sections are parasagittal, left is anterior and up is dorsal. All scale bars 500 µm.

## Thalamic nucleus of the song system DLM is adjacent to but separate from cerebellar-recipient regions of dorsal thalamus

In addition to mapping DT_CbN_, we used similar methods to define the borders of thalamic song system nucleus DLM, so that we could show whether injections of lentiviral vector in either region of dorsal thalamus resulted in labeled axon terminals within Area X. We mapped DLM by injecting tracer in Area X (Fig. 8), since the main target of Area X projection neurons is DLM (Luo & Perkel, 1999a, 1999b). Injections of AAV-GFP in Area X (Fig. 8A) produced anterograde label of the calyceal terminals formed by projection neurons of Area X in DLM (Figure 8B-D). We noticed that anterograde label from Area X injections was always confined to an area of dorsal thalamus that appeared as a bright oval when viewed with a DIC objective, presumably due to heavier myelination (Fig.8B-D). Based on this observation, we concluded that this bright area can serve as a marker for the borders of DLM across animals. We never saw anterograde label within DLM, as defined by this brighter oval, from injections in CbL. We cannot rule out the possibility that CbI or CbM might project to DLM, but as we showed these projections to dorsal thalamus are relatively small compared to the projection from CbL (Fig. 6 and 7). Therefore our results suggest that DLM and DT_CbN_ are two adjacent but distinct areas.

**Figure 8.**
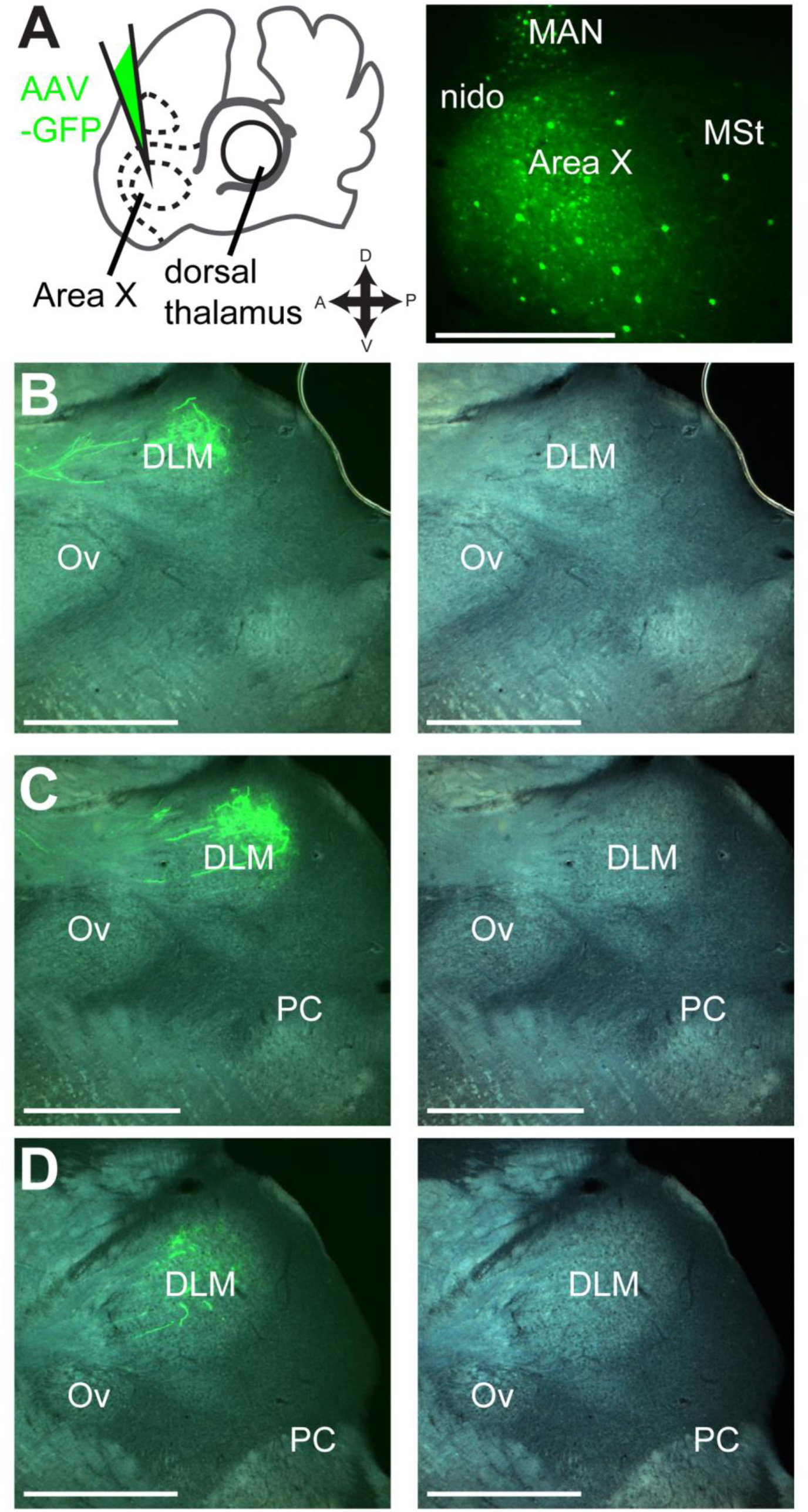
Thalamic song system nucleus DLM is adjacent to but separate from DT_CbN_. **A,** Iontophoretic injection of AAV-GFP vector in Area X. Left panel, schematic of injection site. Right panel, representative section showing injection. nido, nidopallium. MAN, magnocellular nucleus of anterior nidopallium. MSt, medial striatum. **B-D,** Resulting anterograde label of calyceal-like terminals in DLM (dashed white line). Note that label is confined to bright oval as seen in tissue when viewed with DIC filter. DLM, dorsolateral thalamus, medial part. Ov, nucleus ovoidalis. PC, posterior commissure. All sections are parasagittal, left is anterior and up is dorsal. All scale bars: 500 µm

## DLM and immediately-adjacent DT_CbN_ project to Area X

Having produced a map of DLM and DT_CbN_, we next determined whether either region projects to the medial striatum. We made a series of smaller viral injections targeting either DLM or DT_CbN_. For each case, we aligned the injection site with a reference map of DLM and DT_CbN_ (see Methods). We found that whenever the injection site included DLM or the immediately adjacent DT_CbN_ (n=3 hemispheres from two birds, Fig. 9A, E, I), it produced anterograde label in Area X (Fig. 9B-D, F-H, J-L). In two of the three cases, there was strong label of processes with varicosities, similar to what is shown in Fig. 2E, across Area X (Fig. 9D and K, arrow with white outline and no fill). In the remaining case, there was less label in Area X (Fig. 9H, arrow with white outline and no fill) and there was also label of processes with varicosities just ventral to Area X (Fig.9G, solid white arrow). There was no obvious topography in the dorsoventral or anteroposterior planes, although we did often see that label was strongest in the same mediolateral plane as the injection site. Because of space constraints we do not show the entire series here. The entire series from each case shown in Figs. 9 and 10 can be accessed in the Supporting Information (<u>https://doi.org/10.6084/m9.figshare.5437975</u>). We made another set of injections (n=3),which transfected neurons in the more medial and posterior portion of DT_CbN_ while either partially (Fig. 10 A, I) or entirely avoiding DLM (Fig. 10 E). Results from these injections indicated that this more medial and posterior portion of DT_CbN_ does not target Area X. Instead it projects to MSt posterior and ventral to Area X (Fig.10D, F, J, solid white arrows).We did occasionally see small areas of GFP-signal within Area X (Fig. 10J and L, arrow with white outline and no fill) from these injections but strongly-labeled processes with varicosities were posterior and ventral to Area X, and they were much more medial (as in Fig.10F and J). Again label was usually strongest in the same mediolateral plane as the injection site.

**Figure 9.**
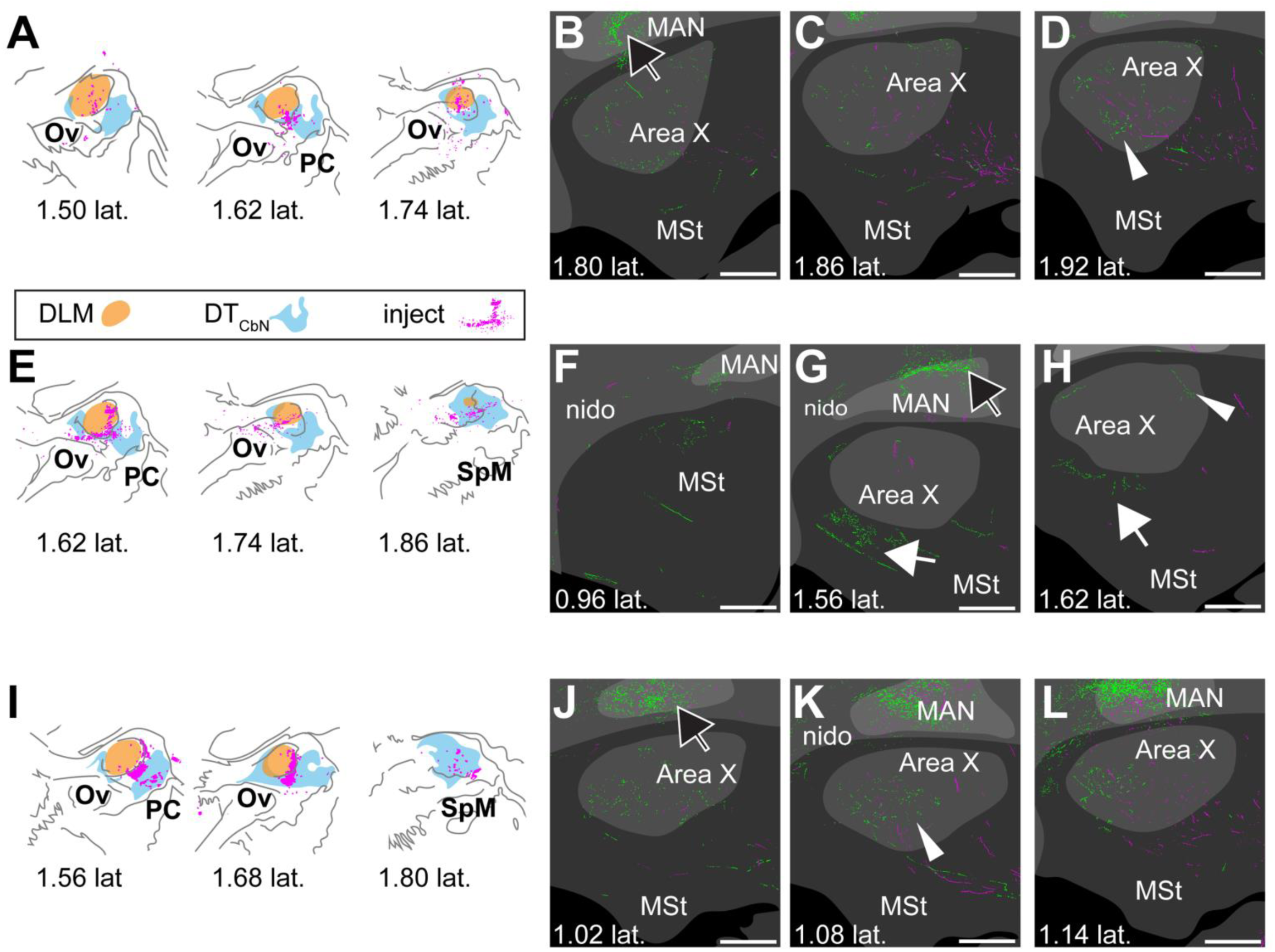
DLM and adjacent DT_CbN_ project to Area X. **A-D,** case where injection was mostly in DLM. Synaptophysin-GFP label was evident in Area X (for example see white arrowhead in **D**) **E-H,** case where injection was in DLM and surrounding DT_CbN_. Synaptophysin-GFP label was evident in medial striatum (**G** and **H**, white arrow) and in Area X (**H**, white arrowhead). **I-L,** case where injection was mostly in DT_CbN._ Although the injection was mainly in DT_CbN_, we again saw strong synaptophysin-GFP label in Area X (for example **K**, white arrowhead). **A,E,I,** Injection sites. Orange region, DLM; cyan region, DT_CbN_; magenta, cell bodies expressing mCherry. **B-D, F-H, J-L.** Series of sections from lateral to medial showing Area X, surrounding medial striatum, and overlying nidopallium. Green, GFP signal. Magenta, mCherry signal. Dark gray, no parvalbumin label; gray, some parvalbumin label; light gray, strong parvalbumin label. White arrowheads, examples of GFP signal in Area X; solid white arrow, GFP signal outside Area X in MSt; black arrow with white outline, GFP signal in nidopallium. DLM, dorsolateral thalamus, medial part. MAN, magnocellular nucleus of anterior nidopallium. MSt, medial striatum. nido, nidopallium. Ov, nucleus ovoidalis. PC, posterior commissure. SpM, medial spiriform nucleus. All sections are parasagittal, left is anterior and up is dorsal. All scale bars 500 µm.

**Figure 10.**
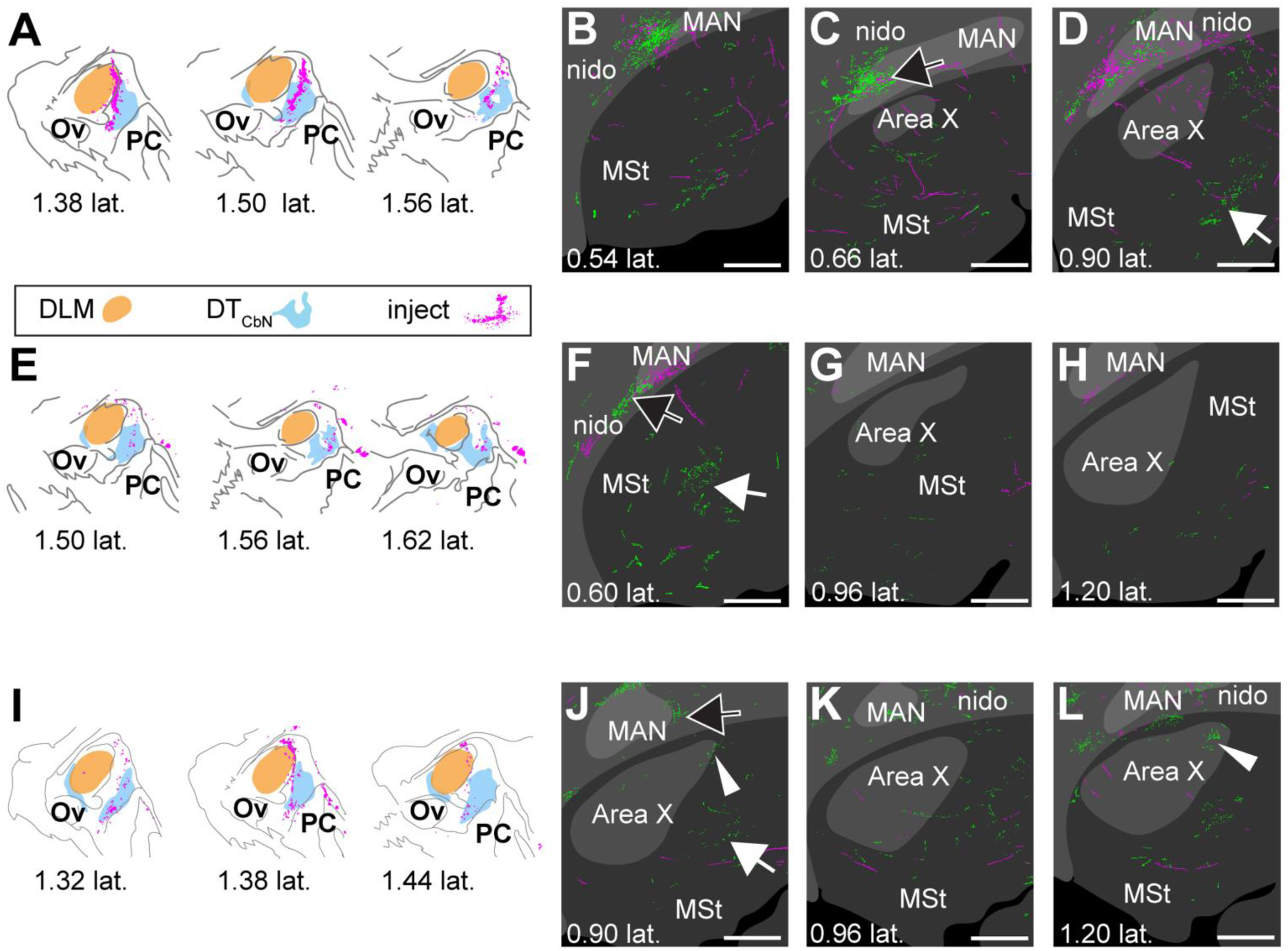
More medial and posterior regions of DT_CbN_ project to medial striatum. **A-D,** case where injection was mostly in DT_CbN_ and more medial. Strong synaptophysin-GFP label was posterior and/or medial of Area X (**D**, white arrow). There was almost no GFP label in Area X in this case. **E-H,** case where injection was in DT_CbN_ but posterior of DLM. Again the strongest Synaptophysin-GFP label was medial of Area X (**F**, white arrow). **I-L,** Area X, arrow with white outline in l. **A,E,I,** Injection sites in DLM and DT_CbN_ for 3 birds. Sections are arranged from medial to lateral reading from left to right. Orange region, DLM; cyan region, DT_CbN_; magenta, cell bodies expressing mCherry. **B-D, F-H, J-L,** series of sections from lateral to medial showing Area X, surrounding medial striatum, and overlying nidopallium. Green, GFP signal. Magenta, mCherry signal. Dark gray, no parvalbumin label; gray, some parvalbumin label; light gray, strong parvalbumin label. Arrow with white outline and no fill, GFP signal in Area X; solid white arrow, GFP signal outside Area X in MSt; black arrow with white outline, GFP signal in nidopallium. DLM, dorsolateral thalamus, medial part. MAN, magnocellular nucleus of anterior nidopallium. MSt, medial striatum. nido, nidopallium. Ov, nucleus ovoidalis. PC, posterior commissure. SpM, medial spiriform nucleus. All scale bars 500 µm.

Regardless of injection site we saw strong label of synaptophysin-GFP and mCherry processes in nidopallium, the cortical layer overlaying the basal ganglia. Injections that included DLM produced label in cortical song system nucleus LMAN as expected, given it is the known target of DLM (for example Fig. 9G and J, black arrow with white outline), and injections in DT_CbN_ separate from DLM tended to produce label in nidopallium outside of LMAN but still within an area of somewhat stronger labeling for parvalbumin (Fig. 10C,F, and J, black arrow with white outline). Since the injections produced label both in cortex as just described, and in the basal ganglia, these results suggest that either individual neurons in dorsal thalamus project to both striatum and cortex, or that thalamic neurons that project to striatum are intermingled with those that project to cortex.

## Discussion

The main result of this paper is that, in Bengalese finches, the basal ganglia nucleus Area X of the song system receives input from two regions of dorsal thalamus: the thalamic nucleus of the anterior forebrain pathway, DLM, and immediately-adjacent regions of cerebellar-recipient thalamus, which we refer to as DT_CbN_ (Fig. 11). Below we address technical considerations and relate our results to previous work. Then we discuss possible functions of the pathways we identified and how future studies might test for these functions.

**Figure 11.**
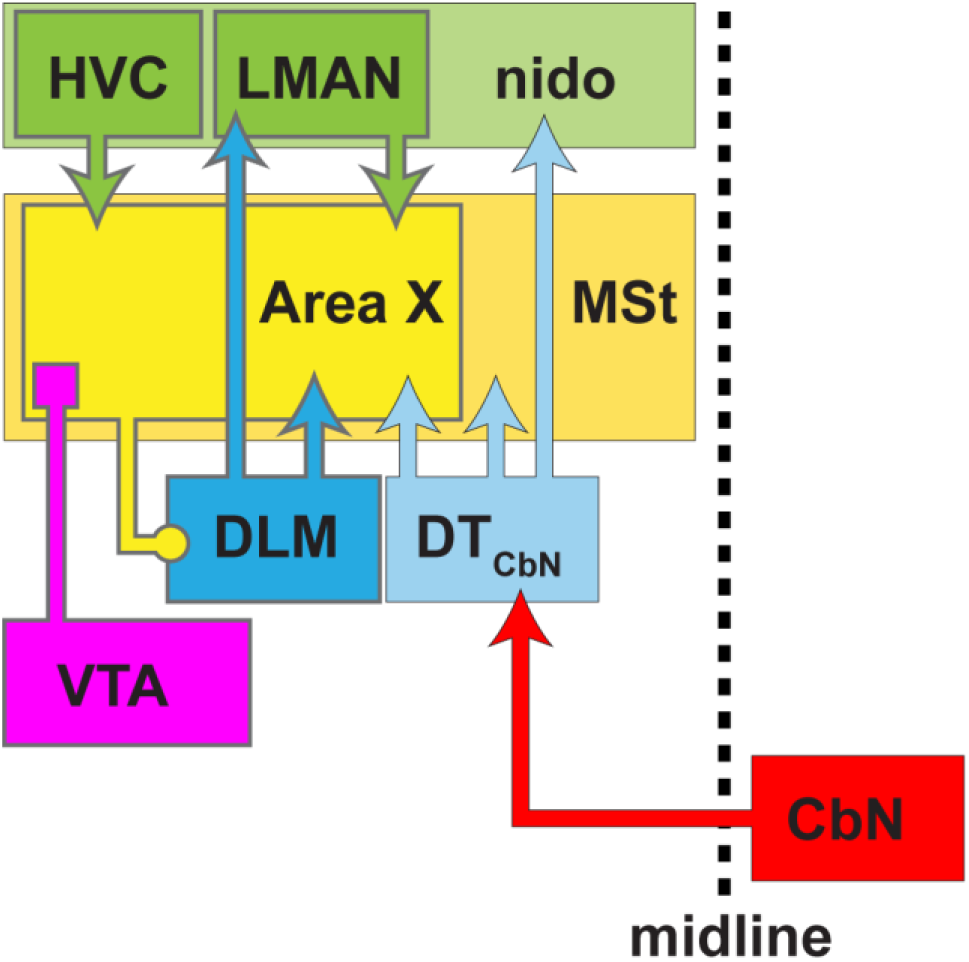
Summary of results. We showed that, in the Anterior Forebrain Pathway (AFP) of the song system, the basal ganglia nucleus Area X receives input from the thalamic nucleus, DLM, as well as adjacent subregions of cerebellar-recipient dorsal thalamus (DT_CbN_) (blue arrows). Hence DLM provides feedback to Area X similar to projections from thalamus to the striatum in mammals, and DT_CbN_ provides a route for output from the cerebellar nuclei (CbN, red arrow) to reach the basal ganglia in the song system through thalamus. More posterior and medial regions of DT_CbN_ project to medial striatum outside of Area X (lighter blue arrow). We also found that DT_CbN_ projects to nidopallium outside of cortical song system nucleus LMAN (lighter blue arrow), implying that DT_CbN_ may communicate both with the song system and with general motor areas outside the song system. Canonical nuclei of the song system are outlined in gray.

## Thalamostriatal projections

We provide strong evidence that dorsal thalamus projects to the medial striatum in a songbird (Fig. 2, Fig. 3). Although this was suggested by retrograde label in prior anatomical studies (Castelino, Diekamp, & Ball, 2007; Lewis, Ryan, Arnold, & Butcher, 1981; Person et al., 2008; Veenman, Karle, Anderson & Reiner, 1995; J. Wild, 1987), several methodological confounds have prevented a definitive demonstration, as recognized previously (Bottjer et al., 1989; Gale & Perkel, 2010; Person et al., 2008). Briefly, the confounds are:(1) passing fibers in the basal ganglia en route from thalamus to cortex could pick up tracer that would retrogradely label thalamic neurons, and this would be indistinguishable from retrograde label due to actual thalomstriatal synapses; (2) Area X and surrounding medial striatum project directly to dorsal thalamus, and so it is not clear whether label in Area X from standard tracers injected in dorsal thalamus has traveled anterogradely or retrogradely. Therefore, to demonstrate whether these projections exist, a method is needed to anterogradely label presynaptic segments of axons of thalamic neurons. The lentiviral vectors we used infected the cell body, yielding signal that only traveled in the anterograde direction (Grinevich et al., 2005; Roberts et al., 2008), and we specifically labeled presynaptic terminals in the basal ganglia using a vector encoding synaptophysin tagged with GFP (Fig. 2, Fig. 3). The synaptophysin-GFP labeled processes had varicosities suggestive of *en passant* synapses (Fig. 2, Fig. 3), as reported for the thalamostriatal system in mammals (Berendse & Groenewegen, 1990; Deschenes, Bourassa, & Parent, 1996). There were also mCherry-labeled axons with the same morphology, implying that these processes occur naturally and did not arise because of the ectopic expression of synaptophysin (Fig. 2). The approach we have taken here therefore complements previous reports of possible thalamostriatal and addresses confounds faced by those previous reports. Another approach, commonly used to identify thalamostriatal synapses in mammals, is to stain for glutamate receptors vGlut1 and vGlut2, since it is known that thalamostriatal axon terminals in mammals preferentially express vGlut2 (while corticostriatal terminals express vGlut1) (D. V. Raju, Shah, Wright, Hall, & Smith, 2006; Dinesh V Raju & Smith, 2005). Future studies could test whether this approach would work in songbirds, as suggested previously (Gale & Perkel, 2010), although we note that to our knowledge only one antibody is available for vGlut2 in birds, and current reports suggest it would not strongly label the basal ganglia (Karim, Pervin, & Atoji, 2015; Karim, Saito, & Atoji, 2014).

## Cerebellothalamic projections

We carried out detailed studies of the cerebellothalamic projection in a songbird, presenting a map of the regions of dorsal thalamus targeted by CbL in Bengalese finches (Fig. 4). Notably, anterograde label from injections in CbL was always outside of song system thalamic nucleus DLM (Fig. 4, Fig. 5, Fig. 8), as reported previously for zebra finches (Person et al., 2008), although there were heavily-labeled regions immediately posterior to DLM (Fig. 4, Fig. 5). We noted that in some cases there was label in dorsal thalamus due to “retrograde” label of collaterals from SpM (H. Karten & Finger, 1976) but based on the very sparse label seen ipsilaterally even when SpM was strongly labeled (see Supporting Information), we are confident the majority of label traveled anterogradely from CbL. To confirm our results showing anterograde label in DT_CbN_, we made injections in dorsal thalamus and showed retrograde label of CbL, consistent with previous work (Vates et al., 1997). We also demonstrated retrograde label in CbI and CbM (Fig.6, Fig.7), a finding that as far as we know has not been been reported previously for songbirds, although it has been reported that neurons in all regions of CbN project to to thalamus in pigeons (Korzeniewska & Güntürkün, 1990; L. Medina & Reiner, 1997; J Martin Wild, 1988; Wylie, Glover, & Lau, 1998) and mammals (Asanuma, Thach, & Jones, 1983; Haroian, Massopust, & Young, 1978; Hoshi et al., 2005; Sugimoto, Mizuno, & Itoh, 1981; Tracey, Asanuma, Jones, & Porter, 1980).

## Projections to Area X from DLM and DT_CbN_

We mapped DT_CbN_ as well as DLM so that we could determine which region of dorsal thalamus gives rise to projections to Area X. We were surprised to find that both DLM and a subregion of DT_CbN_ adjacent to DLM project to Area X (Fig. 9). We do not think this finding can be explained by spill of viral vector from DLM into DT_CbN_ or vice versa; note that we saw strong synaptophysin-GFP label in Area X when the injection was mostly contained within DLM (Fig.9A) and when it was mostly contained to DT_CbN_(Fig.9I). We also note that injection sites in dorsal thalamus as defined by mCherry signal were not noticeably different from sites defined by synaptophysin-GFP signal (which did not fill cell bodies enough to image with the widefield microscope used to record injection sites, but was obvious when imaged with a confocal). Future studies could expand on our results using trans-synaptic tracers, although our understanding is that such methods do not currently work in songbirds (Mundell et al., 2015).

## Functional considerations

One proposed function of thalamostriatal projection in mammals is to convey surprising stimuli that can “rebias” ongoing action selection (Bradfield, Hart, & Balleine, 2013; Minamimoto, Hori, & Kimura, 2009; Smith et al., 2014). Another proposed function is to provide a motor efference copy from lower motor centers (Fee, 2012). However for DLM to perform either of these functions it would require ascending inputs from lower brain areas, which have not been reported. Current models of the songbird AFP do not include a thalamostriatal projection (Fee & Goldberg, 2011), but they propose that a function of the AFP is to modulate the variability of song; the models posit that the brain uses this variability to improve and maintain the ability of the bird to sing (Dhawale, Smith, & Ölveczky, 2017; Fee & Goldberg, 2011; Kuebrich & Sober, 2015; S. Woolley & Kao, 2015). With respect to these models, an alternate hypothesis for the function of the projection from DLM to Area X would be that *en passant* synapses that DLM neurons form with multiple neurons in Area X allow DLM to contribute to correlated activity across neurons in the AFP. If for example single DLM neurons contact multiple medium spiny neurons (MSNs) in Area X, similar to the way that thalamic neurons contact multiple MSNs in the mammalian striatum (Deschenes, Bourassa, Doan, & Parent, 1996; Deschenes, Bourassa, & Parent, 1996; Kuramoto et al., 2009; McFarland & Haber, 2001; Parent & Parent, 2005), this would position them to correlate activity across the multiple neurons with which they form synapses. Modeling studies suggest correlations in activity enable neural networks to produce behavioral variability such as that generated by the AFP (Darshan, Wood, Peters, Leblois, & Hansel, 2017). One way to test for this function would be to reversibly and specifically inactivate thalamostriatal synapses in Area X and measure the effects on vocal variability.

Physiological studies would also be required to address the question of how the cerebellum interacts with the song system via output to dorsal thalamus. A previous physiological study in rats found mixed effects on firing activity of MSNs when optogenetically stimulating regions of CbN that project to thalamus (Chen et al., 2014). Future studies in songbirds should determine at the physiological level how input from the cerebellum via DT_CbN_ modulates activity of target neurons in Area X. Work in mammals suggests the most likely target would be MSNs (Dube, Smith, & Bolam, 1988) and cholinergic interneurons (Lapper & Bolam, 1992)—both cell types are found in Area X (Carrillo & Doupe, 2004; Reiner et al., 2004).

Although we showed that a subregion of DT_CbN_ projects to Area X, our other results on the projections of CbL and DT_CbN_ raise questions about what the function of the cerebellum’s interactions with the song system might be. We found, as previous papers have (Arends & Zeigler, 1991; Person et al., 2008), that CbL targets not just DT_CbN_ but also many sites throughout the midbrain (see Appendices). Because CbL targets these other areas, it seems very unlikely that there is a song-system specific region of CbN. It could be possible that there are single neurons in CbL that synapse only with neurons in DT_CbN_ projecting to Area X, although in mammals it is thought that single neurons in CbN that project to thalamus also send collaterals to the red nucleus (Shinoda, Futami, Mitoma, & Yokota, 1988). Based on the other regions it targets, it is likely that CbL interacts with previously described descending pathways in the avian brain (J. Wild & Williams, 2000; J. M. Wild, 1992) with connections similar to the pyramidal tract in mammals. To our knowledge, the connections of CbL with these pyramidal tract-like pathways in birds have not been studied behaviorally, and even in mammals the functions of these pathways remain an open question (Horne & Butler, 1995; Houk, Keifer, & Barto, 1993; Shmuelof & Krakauer, 2011). Our results also raise the possiblity that the same subregion of DT_CbN_ that projects to Area X also projects to nidopallium just outside of LMAN (Fig. 9, Fig. 10). Future studies should test this using some technique that allows labeling single neurons (Deschenes, Bourassa, Doan, et al., 1996; Kuramoto et al., 2009). If this were the case, it could be that the function of these projections is to help co-ordinates neural activity in the song system with activity in motor systems that control muscles involved in both song and non-song behaviors. By the same token, we emphasize that we also showed that the more medial and posterior part of DT_CbN_ projects to medial striatum outside of Area X (Fig. 9). Assuming that the song system evolved from already existing motor pathways (Farries, 2001; Feenders et al., 2008), then, our findings imply that a pathway from the cerebellum to the basal ganglia through thalamus may have existed before the song system evolved. It remains unclear if these projections contribute directly to motor control in general or if they have some other function such as task-level control of behavior (Bradfield et al., 2013; DeLong & Wichmann, 2009; Minamimoto et al., 2009; Smith et al., 2014) but our results suggest these pathways may be a general feature of motor systems in vertebrate brains.

## Data accessibility

<u>https://doi.org/10.6084/m9.figshare.5437975</u>

## Acknowledgements

This work was supported by NIH grants R01NS084844 and R01DC014364 and NSF grants IOS-1456912, IOS-1457206, and IOS-1451034. Imaging acquisition was supported by the Emory University Integrated Cellular Imaging Microscopy Core of the Emory Neuroscience NINDS Core Facilities grant, 5P30NS055077. We are grateful to the Maney and Jaeger labs at Emory University for material support.

## Notes

**Conflict of Interest Statement:** The authors declare no conflicts of interest.

